# miR-210 is essential to retinal homeostasis in fruit flies and mice

**DOI:** 10.1101/2024.01.30.577988

**Authors:** Davide Colaianni, Federico Virga, Annamaria Tisi, Chiara Stefanelli, Germana Zaccagnini, Paola Cusumano, Gabriele Sales, Mircea Ivan, Fabio Martelli, Daniela Taverna, Massimiliano Mazzone, Cristiano Bertolucci, Rita Maccarone, Cristiano De Pittà

## Abstract

miR-210 is one of the most evolutionarily conserved microRNAs. Recent studies in *Drosophila melanogaster* have unveiled that the absence of miR-210 leads to a progressive retinal degeneration characterized by the accumulation of lipid droplets and disruptions in lipid metabolism. Further investigation into lipid anabolism and catabolism revealed significant alterations in gene expression within these pathways. We provide the first morphological characterization of miR-210 KO mice retinas, highlighting a significant photoreceptor degeneration. While exploring potential parallels between miR-210 KO models in flies and mice, we examined mice lipid metabolism, circadian behaviour, and retinal transcriptome yet found no resemblances, suggesting divergent mechanisms of retinal degeneration between the two species. Simultaneously, analysis of the transcriptome in the brains of miR-210 KO flies revealed the potential existence of a shared upstream mechanism contributing to retinal degeneration in both fruit flies and mammals.

## Introduction

MicroRNAs (miRNAs) are small non-coding RNAs of 21-25 nucleotides in length which negatively regulate protein-coding gene expression at post-transcriptional level by targeting mRNAs and triggering either translational repression or mRNA degradation^1^. Since miRNAs are estimated to regulate approximately 60% of all protein-coding genes and since each miRNA can target up to several hundred mRNAs^2^, it is not surprising their prominent role in a wide range of different biological processes, both under physiological^3^ and pathological conditions^4^. In fact, miRNAs have been implicated in development, differentiation, and proliferation, as well as metabolism, immune responses, and apoptosis^3^. MiRNAs mutations, dysregulation or dysfunction have been linked to the development and the progression of a variety of diseases, including cancer, viral and immune-related diseases, and neurodegenerations^4^. miR-210 (miR-210-3p) is one of the most evolutionarily conserved microRNAs, featuring a “seed sequence” that exhibits 100% identity across flies, mice, and humans^5^. In humans, miR-210 plays a pivotal role in a wide array of biological processes, encompassing cell proliferation, differentiation, stem cell survival, mitochondrial metabolism, angiogenesis, neurogenesis, immune system regulation, DNA repair, apoptosis, and notably, it is a key player in the response to hypoxia^6,7,8,9^. Hypoxia often serves as a trigger for the aforementioned cellular processes^7^. Hypoxia-inducible factors (HIFs), among the most sensitive physiological detectors of hypoxia, orchestrate the expression of a cascade of downstream genes responsible for cell and tissue responses to oxygen deficiency. This regulatory process includes the up-regulation of several specific hypoxia-inducible microRNAs, often referred to as “hypoxamiRs”, with miR-210 being the foremost among them^6,7^. Accordingly, miR-210 has been shown to play a role in many hypoxia-related diseases^10,11^, particularly in cardiovascular diseases^7,12^ and cancer^8,11,13^. Nevertheless, miR-210 is not merely a passive participant in hypoxia^8^, and it continues to unveil novel roles and molecular functions, some of which are unrelated to hypoxic conditions. Interestingly, the fruit fly *Drosophila melanogaster* has emerged as a valuable model for investigating the physiopathological effects of miR-210 dysregulation. Despite the strong conservation of the HIFs pathway between mammals and flies^14^, it appears that the molecular function of miR-210 in response to hypoxia is not preserved^5^. In the same study, Weigelt and colleagues reported that the loss of miR-210 in fruit flies resulted in a progressive retinal degeneration^5^. More specifically, they showed an altered arrangement and morphology of photoreceptor cells, with a progressive decline leading to a complete disruption of the ommatidium structure, also accompanied by a reduction in photoreceptor potential^5^. Afterwards, Lyu and colleagues highlighted the presence of abundant lipid droplet structures within the pigment cells of the retina of miR-210 knock-out (KO) flies^15^, which might represent the cause of the retinal degeneration. In addition, they reported some alterations in lipid metabolism, with increased levels of triacylglycerols (the major storage lipids of lipid droplets) and decreased levels of diacylglycerols^15^, that could only be partially attributed to the role played by the Sterol Regulatory Element-Binding Protein (SREBP)^16^, whose mature and active form was found to be elevated in miR-210 KO flies^15^. Previously, other studies had reported a dysregulation in miR-210 expression levels in specific eye diseases, particularly in proliferative retinopathies, in both mice^17^ and humans^18^. However, the association between miR-210 and these diseases has primarily been established based on its roles in hypoxia response and angiogenesis^19^. Conversely, the research conducted by Weigelt^5^ and Lyu^15^ on miR-210 KO flies has raised the prospect of one or more roles for miR-210 in the eye’s physiology and the visual system that are distinct from the previously documented functions. Interestingly, even miR-210 overexpression has been shown to induce visual impairments^20^. Specifically, when miR-210 was overexpressed in clock cells during the development of flies, it resulted in an altered morphology of the large ventral lateral neurons (l-LNvs) cell bodies and in aberrant arborisations within the optic lobes, which were in turn associated with visual defects^20^. Recent studies have further supported the notion that miR-210 plays distinct roles in maintaining the proper homeostasis of the eye, influencing the cornea^21^ and the trabecular meshwork^22^. This study delves into the morphological characterization of retinas from miR-210 KO mice^23^ using confocal immunofluorescence and transmission electron microscopy, coupled with gene expression analysis. This exploration aims to investigate the potential conservation of miR-210’s role in flies. Despite identifying conserved photoreceptor degeneration between miR-210 KO flies and mice, our attempt to uncover similarities in lipid metabolism or circadian behaviour between these models yielded no conclusive findings. These results imply that distinct mechanisms might drive retinal degeneration in mice, independent of those observed in *Drosophila*, possibly involving species-specific post-transcriptional pathways governed by miR-210. Furthermore, we investigated the outcomes from RNA-seq experiments conducted on miR-210 KO mice retinas and miR-210 KO flies’ brains, exploring potential mechanisms underlying the observed retinal degeneration phenotypes.

## Results

### Loss of miR-210 leads to retinal degeneration and alterations in lipid metabolism in *D. melanogaster*

To deepen the role of miR-210 (*dme-miR-210-3p*) in fruit fly retina, we conducted additional characterization of the miR-210 knock-out (KO) model and explored the visual system of fruit flies overexpressing miR-210. miR-210 expression levels in both miR-210 KO and overexpressing flies, as well as in their relative controls, are reported in Figure S1. We also confirmed, as previously reported by Weigelt and colleagues^5^, that miR-210 is characterized by a tissue-specific expression, with high levels in the fly head/brain and lower levels in the muscles and in the fat body (Figure S2). *D. melanogaster* compound eye is composed by ∼800 independent unit eyes called ommatidia, and each ommatidium includes eight photoreceptors (R1-R8); however, since R7 is localized above R8, only seven photoreceptors are visible in the ommatidium cross-section^37^. Firstly, we examined the ommatidial structure in the retina of 5-day-old flies using transmission electron microscopy (TEM). As previously reported, miR-210 KO flies showed an ongoing photoreceptor degeneration (Figure 1A-B), with an abnormal arrangement of the ommatidial structure and an altered morphology of the photoreceptor neurons, as well as a number of vacuoles that Lyu and colleagues^15^, but not Weigelt and colleagues^5^, identified as lipid droplets. On the other hand, the ommatidia of flies overexpressing miR-210 in retinal cells did not show any defects when compared to relative controls (Figure 1C-D).

**Figure 1.**
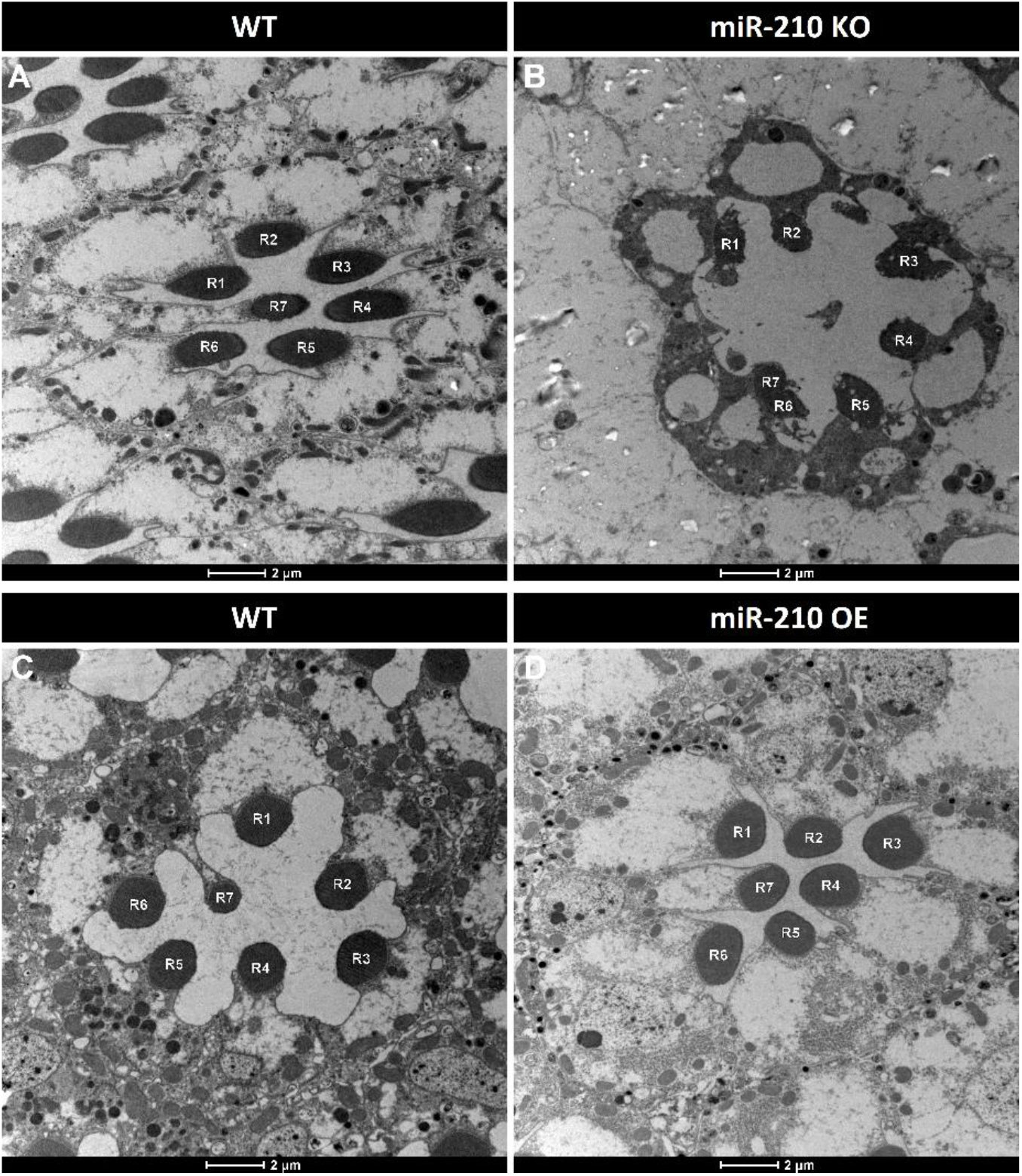
Transmission electron microscopy (TEM) analysis of ommatidial structure in the retina of miR-210 KO and OE flies. Ommatidium cross-section of the retina of 5-day-old flies. When compared to controls (**A**), miR-210 KO flies (**B**) showed aberrant ommatidial structure, with an altered morphology of the photoreceptor neurons indicating an ongoing retinal degeneration. On the other hand, when compared to their relative controls (**C**), flies overexpressing miR-210 in retinal cells (**D**) did not show any ommatidial structural alteration. Scale bar: 2 μm. R1-R7 = photoreceptor neurons. Each image is representative of at least three independent samples.

Based on previous results obtained by Lyu *et al*. (2021)^15^, we further characterized the lipid metabolism, and particularly the triacylglycerol metabolism occurring in the retina of miR-210 KO fly model. The expression levels of the main genes involved in both lipid anabolism (Figure 2A) and lipid catabolism (Figure 2D) pathways^16^ were measured in the heads of 5-day-old miR-210 KO flies and relative controls by qRT-PCR. In the heads of miR-210 KO flies, Lyu *et al*. (2021) observed elevated levels of the mature and active form of sterol regulatory element-binding protein (m-SREBP)^15^. This m-SREBP plays a crucial role in regulating the transcription of genes involved in *de novo* lipogenesis, specifically *Acetyl-CoA-carboxylase* (*ACC*) and *Fatty acid synthase* (*FASN*), both of which were found to be upregulated in miR-210 KO flies^15^. To understand this phenomenon, we measured the expression levels of *SREBP*, of the lipid-sensing chaperone *SREBP cleavage-activating protein* (*SCAP*), and of the *site-specific proteases 1 and 2* (*S1P* and *S2P*), which collectively control SREBP’s nuclear translocation and subsequent activation (Figure 2A, highlighted in green). Surprisingly, in miR-210 KO flies, the expression of *SREBP* and *S1P* was significantly reduced when compared to the control group, indicating a general downregulation in the SREBP activation pathway (Figure 2B). The fatty acids, either from the diet or synthesized through *de novo* lipogenesis (Figure 2A, in grey), undergo esterification by glycerol-3-phosphate acyltransferase Gpat4 (and/or Mino) and lysophosphatidic acid acyltransferase Agpat3^16^. Subsequently, the resulting phosphatidic acid is converted to diacylglycerol by Lipin, further esterified by the diacylglycerol acyltransferase mdy, and stored in lipid droplets (Figure 2A, in blue)^16^. To gain a comprehensive understanding of the lipid metabolism alterations in miR-210 KO flies, we also examined the expression levels of genes encoding these proteins (Figure 2C). We observed a statistically significant down-regulation of *Gpat4* and up-regulation of *mino* in miR-210 KO flies. However, given the potential redundancy in the roles of Gpat4 and Mino^16^, the reported differences in gene expression may not necessarily indicate an alteration in lipid metabolism. Notably, we observed a down-regulation of *Lipin*, a key factor in diacylglycerol synthesis, along with a substantial 2.5-fold upregulation of *mdy*, a crucial player in triacylglycerol production (Figure 2C). These findings are in agreement with and provide a partial explanation for the results obtained by Lyu *et al*. (2021)^15^, whose lipidomic analysis showed a decrease in numerous diacylglycerol species and an increase in various triacylglycerol species in miR-210 KO flies.

**Figure 2.**
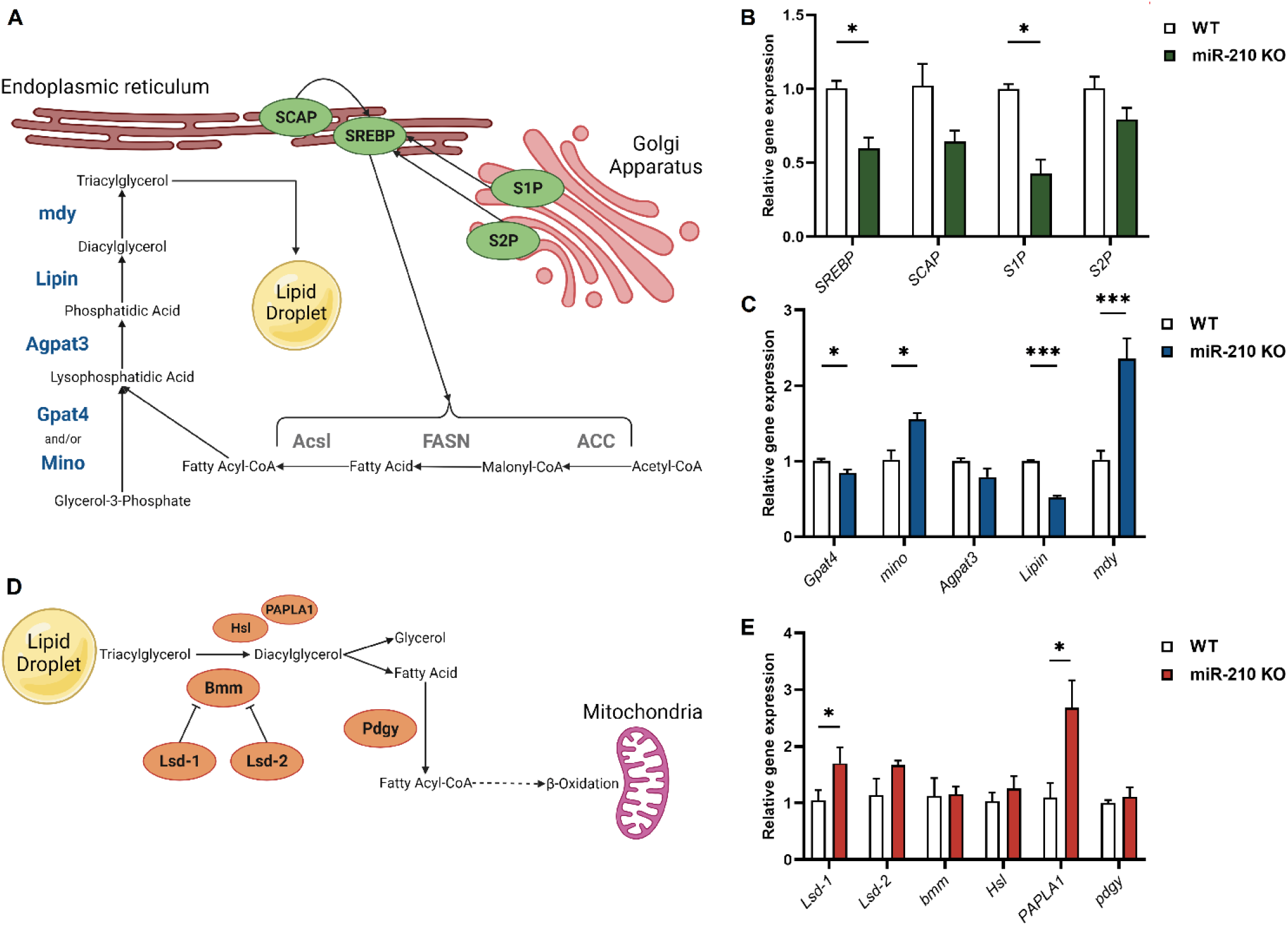
Gene expression analysis of the main genes involved in lipid anabolism and catabolism in miR-210 KO flies. (**A**) A schematic overview of the main proteins involved in lipid anabolism: the SREBP activation pathway, leading to the synthesis of proteins involved in the *de novo* lipogenesis (reported in grey), is reported in green, while the enzymatic synthesis of triacylglycerols is reported in blue. (Created with BioRender.com) (**B**) qRT-PCR expression levels of the main genes involved in the SREBP activation pathway in the heads of 5-day-old miR-210 KO flies and controls. (**C**) qRT-PCR expression levels of the main genes involved in the enzymatic synthesis of triacylglycerols in the heads of 5-day-old miR-210 KO flies and controls. (**D**) A schematic overview of the main proteins involved in lipid catabolism. (Created with BioRender.com). (**E**) qRT-PCR expression levels of the main genes involved in triacylglycerols mobilization and lipolysis in the heads of 5-day-old miR-210 KO flies and controls. The results (N=4) are expressed as mean ± SEM. Student’s t-test was performed to determine significant differences. *p-value < 0.05, ***p-value < 0.005.

We also investigated the expression levels of key genes involved in lipid catabolism. The breakdown of triacylglycerols stored in lipid droplets hinges on the action of lipases. While Brummer (Bmm) is the most well-known lipase in *D. melanogaster*, other enzymes, such as Hormone-sensitive lipase (Hsl) and Phosphatidic Acid Phospholipase A1 (PAPLA1), are thought to participate in the lipolytic process, and there may be undiscovered lipases^16^. Two perilipin-related proteins, Lipid storage droplet 1 and 2 (Lsd-1 and Lsd-2), also known as Plin1 and Plin2, play a role in regulating Bmm activity and lipid droplet catabolism by limiting lipase access to the lipid droplets’ surface^16^. Ultimately, once triacylglycerols are hydrolyzed into diacylglycerol and subsequently into fatty acids, these fatty acids are activated to fatty-acyl-CoA by Pudgy (Pdgy) and then undergo catabolism through β-oxidation (Figure 2D)^16^. In miR-210 KO flies, we observed the upregulation of genes encoding the putative lipase PAPLA1 and, significantly, the lipolysis-regulating enzyme Lsd-1 (Figure 2E). The increased expression of Lsd-1, which restricts Bmm activity, could potentially contribute to the observed pathological phenotype.

In addition, the expression levels of genes showing the higher upregulation (*mdy, PAPLA1*) or downregulation (*SREBP, S1P, Lipin*) in miR-210 KO flies heads were measured also in muscles, in order to investigate whether miR-210 could play a role in lipid metabolism even at the level of muscular tissue. However, we did not observe any differences in gene expression (Figure S3).

Simultaneously, we employed a colorimetric assay to measure the levels of triacylglycerols (TAG) comparing the heads of both starved and non-starved miR-210 KO and WT flies. Surprisingly, we did not observe a difference in TAG content between the two experimental groups (Figure S4). Conversely, when subjected to starvation, we found that the TAG consumption in the heads of miR-210 KO flies was similar, if not increased, compared to that in WT flies (Figure S4). This observation suggests the restoration of normal lipid metabolism in miR-210 KO flies under stressful conditions, indicating that lipid droplet accumulation does not overwhelm the organism’s metabolic demands.

Overall, our research confirms that in miR-210 KO flies, both lipid synthesis (Figure 2A-C) and breakdown (Figure 2D-E) are significantly impaired compared to wild type. These disruptions involve multiple pathways, suggesting that several miR-210 target genes, not exclusively related to lipid metabolism, could be involved, and that lipid droplet accumulation may be a secondary phenotype rather than the primary cause of the observed retinal degeneration.

### The retinal degeneration associated with lack of miR-210 is conserved from flies to mice

Despite the evidence of the induced retinal degeneration in fruit flies, the effects of miR-210 loss in the mammalian retina remained to be investigated. For this purpose, we characterized the retina of miR-210 KO mice (miR-210 expression levels are reported in Figure S5). To evaluate the retinal homeostasis, we performed immunofluorescence on mouse retinal cryosections to detect the neuroinflammation targeting GFAP and Iba-1. It is well known that noxious stimuli induce an inflammatory response of the retinal tissue. This reactivity is detectable looking at the upregulation of GFAP by Müller cells and the microglia activation and migration from the inner retina to the outer nuclear layer^38,39^. In the ventral (inferior) region of the retina (Figure 3A) we found an upregulation of GFAP from the ganglion cell layer (GCL) to the outer nuclear layer (ONL) (Figure 3B-F). Furthermore, a trend of increase in the number of IBA-1-positive (+) cells was detected in the ONL of miR-210 KO mouse retinal cryosections when compared to wild type, associated with a distinct structural alteration in the ONL within the ventral region (Figure 3G-I), likely involving changes in the extracellular matrix. Given the strong indications of ongoing retinal stress from the GFAP and IBA-1 immunofluorescence analyses, we opted to provide a comprehensive morphological characterization of miR-210 KO mouse retinas through TEM. This analysis revealed a clear photoreceptor degeneration affecting the photoreceptor outer segments (OS) of miR-210 KO mice compared to wild type (Figure 3J-K, indicated by the arrows). The higher magnification TEM images better show the loss of ultrastructural morphology of OS of miR-210 KO mice (Figure 3L-M). On the other hand, similarly to fruit flies, the retina of mice overexpressing miR-210 (Figure S6) did not show any alteration when compared to controls (Figure S7). In conclusion, as in *D. melanogaster*, the loss of miR-210, rather than its overexpression, resulted in retinal stress and photoreceptor degeneration in the ventral region of the retina. At the same time, neither the TEM nor the confocal fluorescence microscopy analyses revealed the presence of lipid droplets or any signs of lipid metabolism alterations, which were prominent features of the miR-210 KO fly model.

**Figure 3.**
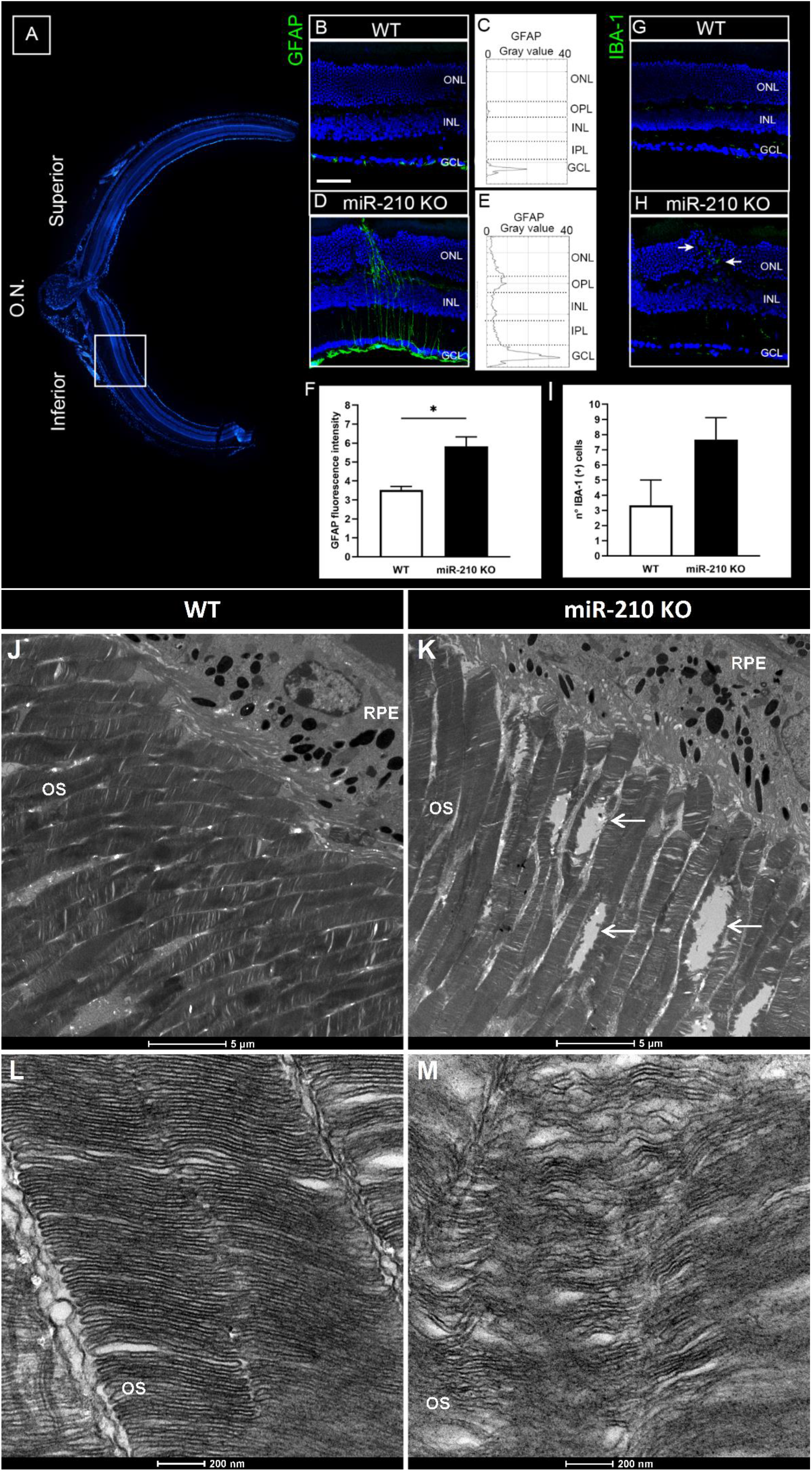
Morphological analysis of the miR-210 KO mouse retina. (**A**) Overview of a retinal section crossing the optic nerve (O.N.) stained with bisbenzimide nuclear dye, showing the superior (dorsal) and inferior (ventral) retina. (**B**-**I**) Analysis of neuroinflammatory markers GFAP (**B**-**F**) and IBA-1 (**G**-**I**). (**B, D**) Representative confocal images of anti-glial fibrillary acidic protein (GFAP) immunostaining acquired at the inferior retina of wild type (WT) (**B**) and miR-210 KO (KO) (**D**) retinal cryosections. Bisbenzimide nuclear dye: blue; GFAP: green. Scale bar: 50 μm. Each image is representative of at least three independent samples. (**C, E**) Profile plots of GFAP fluorescent signals throughout the retinal layers of wild type (WT) (**C**) and miR-210 KO (KO) (**E**) mice. (**F**) Column chart of the fluorescence intensity of GFAP fluorescent signal in the ventral retina of wild type (WT) and miR-210 KO (KO) mice. The results (N=3) are expressed as mean ± SEM. Student’s t-test was performed to determine significant differences. *p-value < 0.05. (**G, H**) Representative anti-ionized calcium binding adaptor molecule 1 (IBA-1) immunostaining acquired at the inferior retina of wild type (WT) (**G**) and miR-210 KO (KO) (**H**) retinal cryosections. The white arrows indicate IBA-1 (+) cells infiltrating in the outer retina at the site of a structural alteration of the outer nuclear layer (ONL). Bisbenzimide nuclear dye: blue; IBA-1: green. Scale bar: 50 μm. Each image is representative of at least three independent samples. (**I**) Column chart of microglia quantification in the outer nuclear layer (ONL) through the entire section from superior to inferior of wild type (WT) and miR-210 KO (KO) retinal cryosections; the measurement is expressed as number of IBA-1 (+) cells in the outer nuclear layer (ONL). The results (N=3) are expressed as mean ± SEM. Student’s t-test was performed to determine significant differences. (**J**-**M**) Transmission electron microscopy (TEM) images showing photoreceptor outer segments (OS) layer (**J, K**) and photoreceptor ultrastructure (**L, M**) in the retina of wild type (WT) (**J, L**) and miR-210 KO (**K, M**) mice. The white arrows indicate the photoreceptor degeneration occurring in the outer segments (OS) of miR-210 KO mice. Scale bars: 5 μm (**J, K**), 200 nm (**L, M**). OS = photoreceptor outer segments; RPE = retinal pigment epithelium. Each image is representative of at least three independent samples.

### Analysis of circadian locomotor activity in miR-210 KO mice

The visual system is essential for the regulation of circadian rhythmicity in mammals, since the endogenous clock constantly needs to be synchronized (*entrained*) by external stimuli (also called *zeitgebers*), and light represents the most powerful of them^40^. Moreover, physical communications between the visual system and the circadian clock have been reported both in *Drosophila melanogaster* and mammals, with several photoreceptors targeting different subsets of clock neurons^40,41^. This close interplay is also reflected at the molecular level: several microRNAs have been shown to exert pleiotropic effects on both visual system and circadian behavior, in particular in *D. melanogaster*^42^. Among them, miR-210 has been reported to have a prominent role in the modulation of circadian outputs in flies. Flies overexpressing miR-210 in clock neurons lost their ability to anticipate the lights-on transition and delayed their evening activity onset under a 12/12 hours light/dark (LD) cycle. At the same time, under constant darkness (DD), miR-210 overexpression led to the disruption of locomotor activity cycles with 70% arrhythmicity^20^. On the other hand, miR-210 KO flies showed an advanced circadian phase under DD and an advanced evening activity onset under LD^20,43^. To explore other potential parallels between miR-210 KO fly and mouse models, we decided to investigate whether miR-210 KO mice exhibited any circadian activity disturbances. We monitored the locomotor activity of miR-210 KO mice and relative controls for 10 days under a 12/12 hours light/dark (LD) cycle and subsequently for 10 days under constant darkness (DD). Different parameters were taken into account: time of activity onset and offset (Figure 4A-D), day-time and night-time activity (Figure 4A-J), periodicity (data not shown), and acrophase (*i*.*e*., the time at which the peak of a rhythm occurs) under LD conditions (Figure 4K). Nevertheless, no circadian alterations were detected between miR-210 KO and WT mice, suggesting that the regulatory influence of miR-210 on circadian rhythms observed in *D. melanogaster* may not be conserved in mammals.

**Figure 4.**
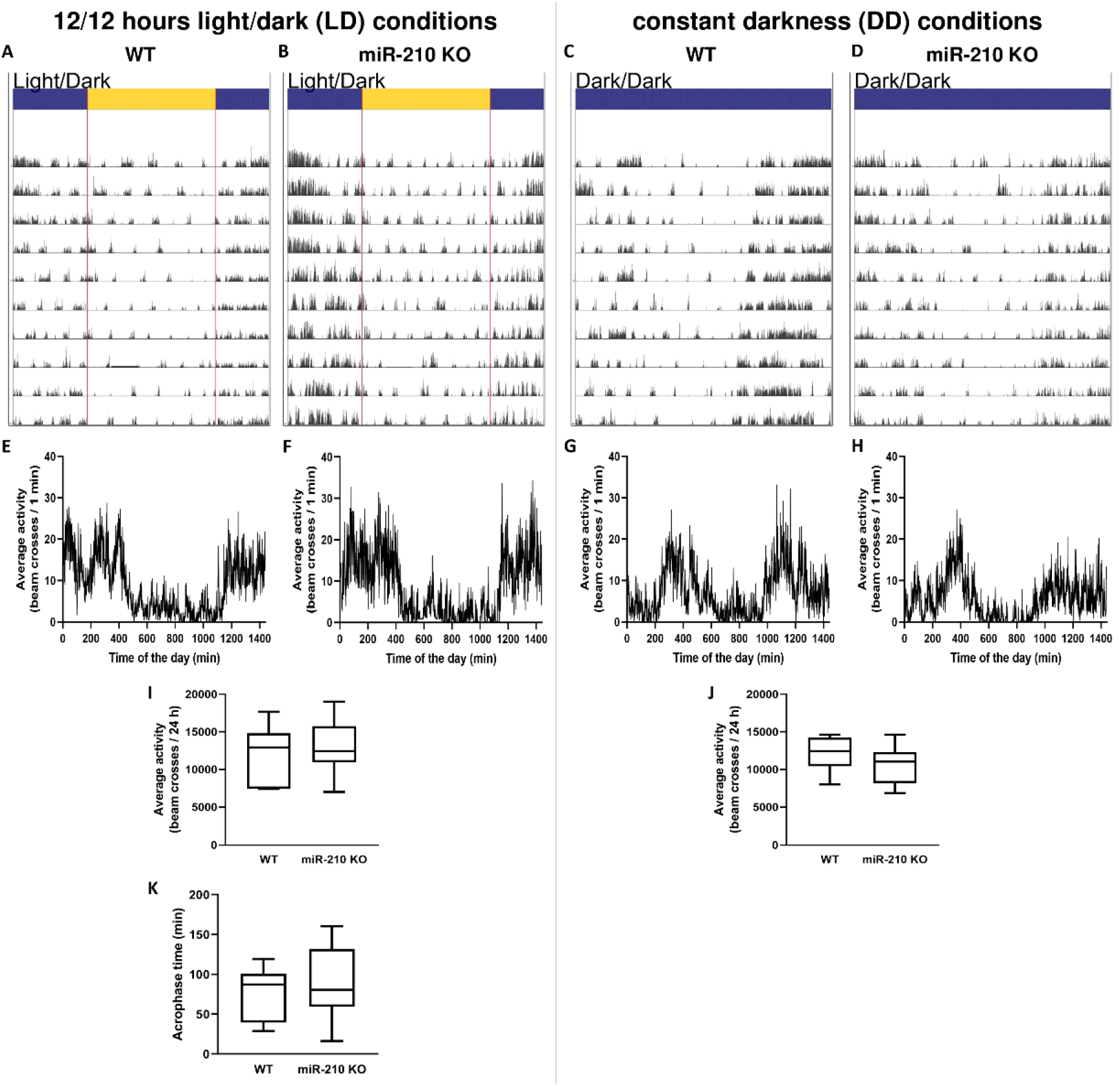
Representative examples of locomotor activity of miR-210 KO mice under LD cycle and constant darkness. The daily and circadian locomotor activity of WT (**A, C**) and miR-210 KO mice (**B, D**) under LD (**A, B**) and DD conditions (**C, D**) is reported as actogram. The corresponding mean waves are plotted in (**E**-**H**). The comparison between WT and miR-210 KO mice locomotor activity under LD (**I**) and DD (**J**) conditions (Student’s t-test), as well as that of the acrophase under LD (**K**) conditions (two-way RM ANOVA), revealed no differences (p-value > 0.05) between the two experimental groups (N=7).

### miR-210 KO mice do not show any alteration in lipid metabolism

As we already mentioned, no lipid droplets, striking features of the miR-210 KO fly model^15^, were detected in the retina of miR-210 KO mouse model. Nonetheless, we sought to investigate the possibility of hidden alterations in lipid metabolism by assessing the expression levels of mouse orthologs of genes involved in lipid metabolism (*Srebf1, Srebf2, Mbtps1, Gpat4, Gpam, Lpin3, Dgat1, Acaca, Acacb, Acss2, Fasn, Acly*) which had been found to be differentially expressed in the heads of miR-210 KO flies. As expected, no significant differences in the retinal expression levels of genes involved in lipid metabolism were detected between miR-210 KO and WT mice (Figure 5). This result, taken together with the absence of lipid droplets accumulation in the retina of miR-210 KO mice, strongly suggests that the molecular mechanism underlying the retinal degeneration is different from flies to mice, or that the alteration in lipid metabolism reported in *D. melanogaster* is secondary to another perturbation that the lack of miR-210 exerts on the cell physiology.

**Figure 5.**
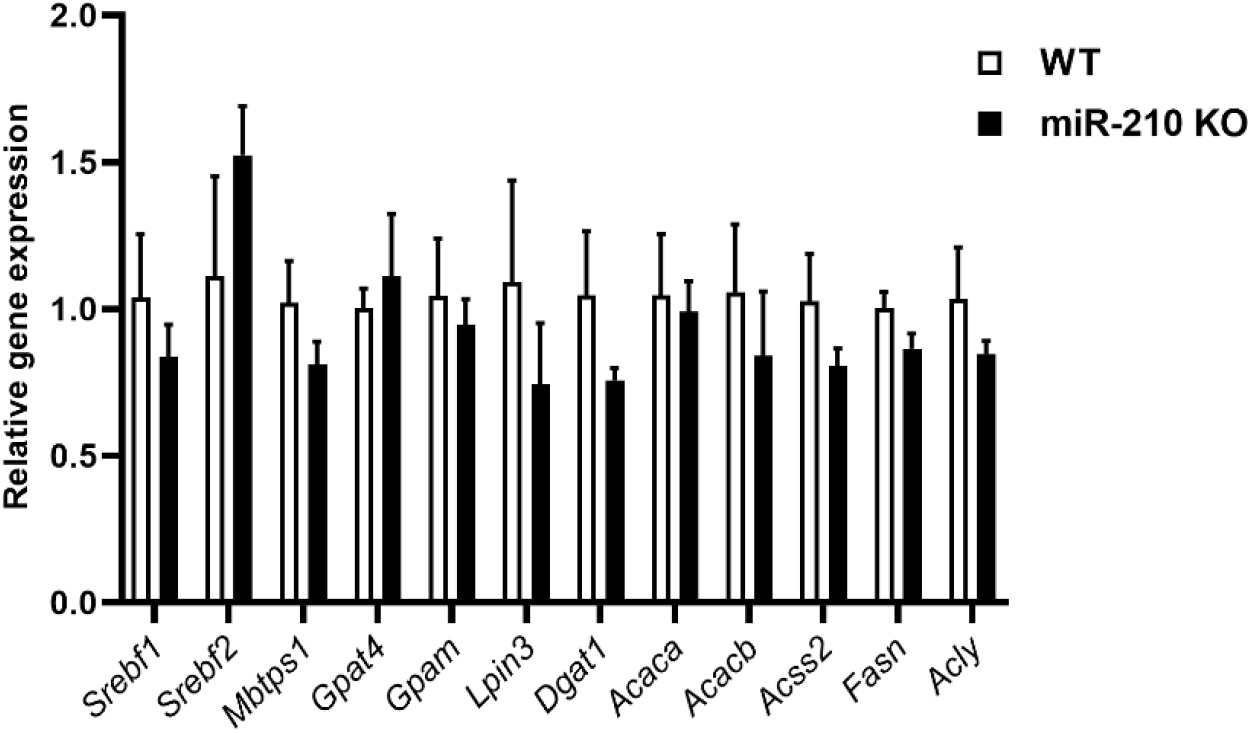
Gene expression analysis of mouse orthologs of genes involved in lipid metabolism in *D. melanogaster*. The expression levels of mouse orthologs corresponding to genes implicated in lipid metabolism, which were identified as differentially expressed in miR-210 KO flies, were evaluated via qRT-PCR in the retinas of miR-210 KO and WT mice aged 10-11 weeks. The results (N=3) are expressed as mean ± SEM. Student’s t-test was performed to determine significant differences.

### Mice retinal degeneration is characterized by altered expression of extracellular matrix genes

Given the unaltered state of lipid metabolism and aiming for a more thorough and comprehensive understanding of the changes taking place in the retinas of miR-210 KO mice, we conducted an RNA sequencing (RNA-seq) analysis. The RNA-seq identified roughly 13,500 genes, of which 107 displayed differential expression. Among these, 38 genes were significantly upregulated, while 69 were significantly downregulated in the retinas of miR-210 KO mice compared to wild type (Table S2). Subsequent gene ontology (GO) analysis (Figure 6, Table S3), as well as reinforcing the lack of involvement of lipid metabolism in the pathological phenotype, also unveiled that the differentially expressed genes were predominantly enriched for cellular components and molecular functions related to chloride channel activity (*Ano1, Cachd1, Clcn1, Gabrb1, Kcng2, Slc1a4, Xntrpc*) (Figure 6B) and, most significantly, with the structure of the extracellular matrix (*Col6a1, Col7a1, Col11a1, Hapln1, Hmcn1, Lama3, Lama5, Umodl1*) (Figure 6C).

### Gene expression analysis of miR-210 KO flies brains reveals neuronal defects in signals detection and transduction possibly attributed to extracellular matrix alterations

Surprisingly, despite the pronounced pathological phenotype in the retinas of miR-210 KO mice (Figure 3), the RNA-seq analysis conducted on miR-210 KO and WT mouse retinas yielded a limited number of DEGs (Table S2), with a majority involved in the maintenance and functionality of the extracellular matrix (Figure 6, Table S3). Moreover, among the genes found to be significantly overexpressed in miR-210 KO mice retinas, we were not able to identify any of the miR-210-3p validated and predicted targets (Table S2). These findings led us to speculate that the observed photoreceptor degeneration may result from upstream neuronal developmental and/or physiological anomalies associated with altered miR-210 expression. Our hypothesis, further described in the next section, is based on the pleiotropic but still poorly characterized effects that miR-210 exerts on the nervous system physiology^44-49^ and pathophysiology^50^, especially when downregulated^48,49^. In *Drosophila*, miR-210 overexpression in clock neurons affects large ventral lateral neurons (l-LNvs) morphology, with altered star-shaped cell bodies and aberrant PDF-positive arborisations in the optic lobes, additionally resulting in visual defects^20^. For these reasons, we decided to perform an RNA sequencing experiment on the brains of miR-210 KO and WT flies. Previous studies by Weigelt^5^ and Lyu^15^ investigated the transcriptome of miR-210 KO flies’ heads, which encompass both the brain and eyes, along with several other tissues to a lesser extent. In both studies, it was observed that the most downregulated genes in miR-210 KO *vs*. WT flies were related to phototransduction and rhabdomere function, while a significant proportion of the upregulated genes were associated with lipid metabolism^5,15^. In our experiment, we were able to detect approximately 9,500 genes, among which 1,053 were differentially expressed (420 were significantly upregulated, while 633 were significantly downregulated) in the brains of miR-210 KO flies with respect to WT controls (Table S4). The GO analysis unveiled that the differentially expressed genes, particularly those that were downregulated, were predominantly enriched in the detection and transduction of light stimuli (Figure 7, Table S5). This finding aligns with the results obtained by Weigelt^5^ and Lyu^15^, suggesting that the absence of miR-210 impacted not only the eyes but also the entire brain. This provides further support for our hypothesis. Moreover, several downregulated genes were involved in processes correlated with actin cytoskeleton and actomyosin structural organization (Table S5), which are essential for the structural integrity of neurons and are required for the generation and plasticity of axons and dendritic spines^51,52^. At the same time, no pathway associated to lipid metabolism was found to be enriched in miR-210 KO flies’ brains (Table S5). Interestingly, among the upregulated genes, the few enriched pathways (all related to *Col4a1, prc*, and *vkg* genes) included “Collagen network”, “Complex of collagen trimers”, and “Collagen type IV trimer”, with the latter serving as a crucial structural element of the basement membrane, a specialized form of the extracellular matrix^53^.

**Figure 6.**
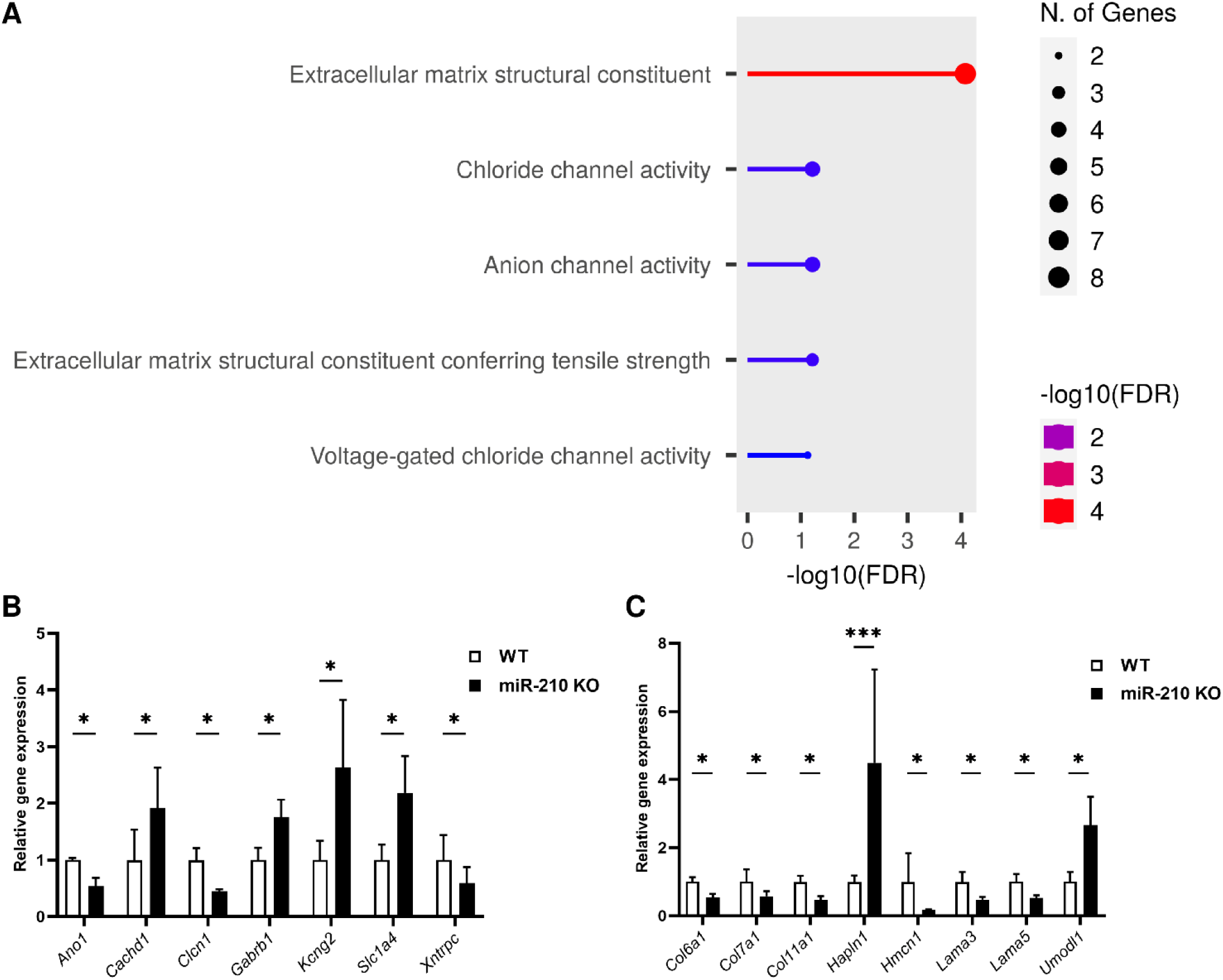
RNA-seq and gene ontology (GO) analysis of differentially expressed genes (DEGs) in the retinas of miR-210 KO mice. **(A)** Gene ontology (GO) analysis was conducted using the ShinyGO annotation tool, focusing on genes exhibiting differential expression between miR-210 KO and WT mice retinas. (**B**-**C**) Differentially expressed genes (DEGs) involved in chloride channel activity and in the maintenance and functionality of the extracellular matrix (**C**), resulting from the RNA sequencing (RNA-seq) analysis.

**Figure 7.**
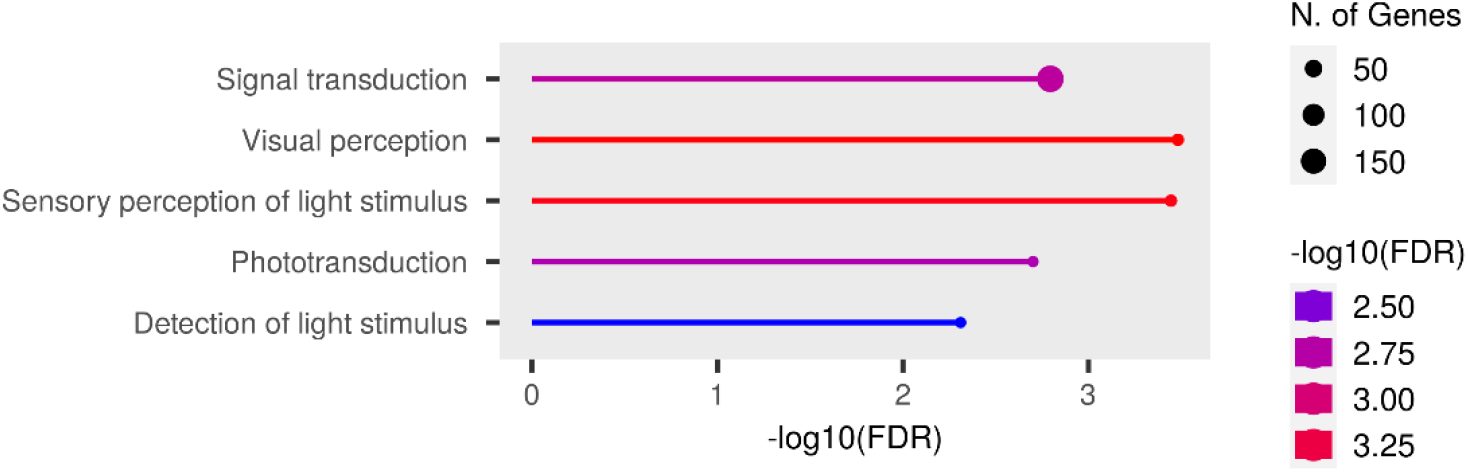
Gene ontology (GO) analysis of differentially expressed genes in the brains of miR-210 KO flies. GO analysis performed using ShinyGO annotation tool starting from the genes found to be differentially expressed between miR-210 KO and WT fly brains. The top five significantly enriched biological processes are depicted.

## Discussion

MicroRNAs (miRNAs) have been involved in a multitude of physiological processes^3^ and pathological conditions^4^ since their discovery. Several miRNAs have been linked to the development and proper function of the eye and the broader visual system, in both *Drosophila melanogaster*^42^ and mammals^54^. Recent evidence suggests that miR-210 is part of this group of miRNAs. miR-210 is one of the most evolutionarily conserved miRNAs, having an identical seed sequence in flies, mice and humans^5^. In mammals, miR-210 is known to be involved in many biological processes, ranging from cell proliferation, differentiation, apoptosis, but especially in the cellular response to hypoxia^6,7^. As a result of its pleiotropic functions and its demonstrated upregulation by hypoxia-inducible factors (HIFs), it has frequently been dubbed “the master hypoxamiR”^7^. However, miR-210 is more than just a silent player in hypoxia^8^, and new molecular functions of this miRNA, not necessarily associated to hypoxic conditions, are being investigated and are emerging. In *Drosophila melanogaster*, although the HIFs pathway is strongly conserved between mammals and flies^14^, the induction of miR-210 expression in response to hypoxia does not seem to be maintained^5^, therefore making the fruit fly an interesting model to study miR-210 roles unrelated to hypoxic conditions. Accordingly, the first evidence of a possible role for miR-210 in the physiology of the eye and of the visual system came from *Drosophila melanogaster*. Two almost contemporary studies, Cusumano *et al*. (2018)^20^ and Weigelt *et al*. (2019)^5^, characterized miR-210 expression pattern, reporting a brain-specific expression, in particular at the level of the retina and of the optic lobes. In addition, Cusumano and colleagues^20^ reported that miR-210 overexpression in clock cells results in an altered morphology of the large ventral lateral neurons (l-LNvs) cell bodies and in aberrant arborisations in the optic lobes, which were in turn associated with visual defects. At the same time, Weigelt and colleagues^5^ reported that miR-210 knock-out (KO) results in a progressive retinal degeneration with a complete disruption of the ommatidium structure. Later, Lyu and colleagues^15^ identified the presence of abundant lipid droplet structures in the retina of miR-210 KO flies as a possible cause of photoreceptors degeneration. They also reported some alterations in lipid metabolism, with increased levels of triacylglycerols and decreased levels of diacylglycerols, only partially explained by the activity of the Sterol Regulatory Element-Binding Protein (SREBP)^16^, whose mature and active form was found to be elevated in miR-210 KO flies^15^. In addition, they performed an RNA-seq experiment on the heads of 5-day-old miR-210 KO and control flies, which showed that most downregulated genes were involved in phototransduction and rhabdomere function while the great part of the upregulated genes were involved in lipid metabolism^15^ (similar results were obtained in an RNA-seq performed by Weigelt and colleagues^5^ on the heads of 1-hour-old miR-210 KO and control flies). Finally, they were able to identify a direct target of miR-210 involved in lipid metabolism, *acetyl-coenzyme A synthetase* (*ACS*), but its reduction in the miR-210 KO background only partially suppressed the retinal degeneration^15^, indicating that other genes might also be implicated in the observed retinal pathological phenotype. In our study, we corroborated the findings of both Weigelt^5^ and Lyu^15^, reinforcing the observation that the loss of miR-210, but not its overexpression, results in retinal degeneration (Figure 1). Subsequently, we concentrated our efforts on investigating alterations in lipid metabolism dissecting both lipid anabolism (SREBP activation pathway, *de novo* lipogenesis, enzymatic synthesis of triacylglycerols) and lipid catabolism (triacylglycerols mobilization and lipolysis) with the aim of identifying changes in the expression of specific genes or pathways (Figure 2A and 2D). Interestingly, even though Lyu *et al*. (2021)^15^ reported a higher presence of the mature and active form of SREBP in the heads of miR-210 KO flies, we observed that the expression levels of the *SREBP* gene, as well as most of the genes constituting the molecular apparatus responsible for regulating SREBP activation (*SCAP, S1P, S2P*)^16^, were downregulated in the heads of miR-210 KO flies compared to the control group (Figure 2B). In addition, we also reported profound alterations in the expression of several other genes, involved in the enzymatic synthesis of triacylglycerols (*Gpat4, mino, Lipin, mdy*) (Figure 2C) as well as in triacylglycerols mobilization and lipolysis (*Lsd-1, PAPLA1*) (Figure 2E). Taken together, this dysregulation across multiple lipid metabolism pathways suggests that it may not be solely a single miR-210 target gene responsible for these changes. Furthermore, these alterations may not be the primary cause of the retinal degeneration, possibly indicating that lipid metabolism disruptions are secondary phenomena. In line with this, our observation that under starvation conditions, the consumption of triacylglycerols in the heads of miR-210 KO flies is similar to, if not higher than what measured in WT flies (Figure S4) suggests that the accumulation of lipid droplets and associated lipid metabolism alterations do not excessively strain the organism’s metabolic requirements. This finding supports the notion that lipid metabolism disturbances are not the primary or sole contributors to the observed retinal pathological phenotype. Afterwards, we investigated whether the repercussions of miR-210 deficiency in the fruit fly retina persisted in mammals. In mice, miR-210 was found to be highly expressed in the adult mice retina^55^, being detected in at least three different layers: the ganglion cell layer (GCL), the inner nuclear layer (INL), and the outer nuclear layer (ONL)^56^. In humans, miR-210 was found in both the retina^57^ and the subretinal fluid^58^, as well as in the vitreous humour^59^, suggesting a possible role in eye homeostasis. In addition, miR-210 has also been associated to some eye diseases since it was found to be upregulated in the retina of a murine model of proliferative retinopathy^17^ and in the serum of diabetic retinopathy^18^ and primary open-angle glaucoma^60^ patients. In the latter, miR-210 expression was associated with visual field defects and average retinal nerve fiber layer thickness^60^. Nonetheless, considering the angiogenetic aspect of these diseases^19,61^, miR-210 was initially hypothesized to be involved primarily due to its established functions in hypoxia response and angiogenetic pathways. To explore the potential preservation of the impact of miR-210 deficiency on the retina from fruit flies to mammals, we conducted a morphological characterization of the miR-210 KO mice retina. This investigation involved employing both confocal immunofluorescence and transmission electron microscopy. The immunostainings using two different neuroinflammatory markers, GFAP and IBA-1, revealed an ongoing retinal stress, particularly at the level of the ventral (inferior) region of the retina, with compromised extracellular matrix structure (Figure 3A-I). This result was further confirmed by the TEM analysis, which revealed a clear photoreceptor degeneration occurring in the retinas of miR-210 KO mice (Figure 3J-M). Notably, neither the confocal fluorescence microscopy nor the TEM analyses revealed the presence of lipid droplets. Conversely, similarly to fruit flies, the retinas of mice overexpressing miR-210 exhibited no detectable alterations (Figure S7). In summary, these findings imply that the retinal degeneration associated with the absence of miR-210 is a conserved phenomenon from flies to mice. Considering the pivotal role of light and, consequently, the visual system in the regulation of circadian rhythms^40^ and given that miR-210 has demonstrated a significant role in modulating circadian processes in flies^20,43^, we explored the possibility of circadian activity alterations in miR-210 KO mice. Nevertheless, our investigations did not reveal any circadian disruptions (Figure 4), indicating that the regulatory function of miR-210 on circadian rhythms observed in *D. melanogaster* is not conserved in mammals. The irrelevant role of miR-210 in the circadian rhythm process in the mammalian retina could be a further demonstration of the functional as well as morphological diversity of the visual system between the two species under study. Based on this evidence, it is possible to corroborate the fundamental role of melanopsin ganglion cells, also known as intrinsically photosensitive retinal ganglion cells (ipRGCs), in the regulation of the circadian rhythm in mammals^62^. We additionally investigated potential alteration in lipid metabolism, striking feature of the miR-210 KO fly model, by assessing the expression levels of the mouse orthologs of genes previously identified as differentially expressed in miR-210 KO flies. However, no similarities were observed between the two miR-210 KO models (Figure 5). Taken together with the absence of lipid droplets accumulation in the retina of miR-210 KO mice, this outcome strongly indicates that the molecular mechanism driving retinal degeneration differs between flies and mice. Alternatively, it suggests that the lipid metabolism disturbances observed in *D. melanogaster* are secondary to other perturbations induced by the absence of miR-210 in cell physiology. In order to have a more comprehensive picture of the alterations occurring in the retina of miR-210 KO mice, we performed an RNA-seq analysis which resulted in a small amount (107) of differentially expressed genes (Table S2), most of which were linked to chloride channels activity and, notably, extracellular matrix structural constituents (Figure 6, Table S3), that are known to play crucial roles in retinal physiology^63-65^. Interestingly, a link between miR-210 and extracellular matrix is recently emerging. Specifically, miR-210 has been observed to impact the expression of type II collagen and aggrecan in nucleus pulposus cells^66^. Furthermore, within the eye, it has been identified as a mediator of trabecular meshwork extracellular matrix accumulation^22^. In ocular contexts, miR-210 has also been found to actively participate in corneal epithelial repair by regulating EphA2/Ephrin-A1 signaling^21^. Notably, the corneal epithelium functions under normoxic conditions, particularly when the eye is open and avascular^21^. This underscores that at least some of miR-210’s roles within the eye are independent of hypoxia. All these evidence indicate direct roles of miR-210 within the eye, and by extension, within the retina. These roles may arise from its potential regulation of chloride channels and extracellular matrix genes, either directly or indirectly. Conversely, another plausible scenario, potentially linking the observed phenotypes in both fruit flies and mice, is that the retinal phenotype may result from an upstream disruption in central nervous system homeostasis. This could partially explain the small number of differentially expressed genes found in the retinas of miR-210 KO mice compared to controls (Table S2). This outcome was somewhat unexpected given the pronounced pathological phenotype. Intriguingly, among the genes found to be overexpressed in miR-210 KO mice retinas, we were unable to pinpoint any of the validated and predicted targets of miR-210-3p (Table S2). Moreover, miR-210 roles in the nervous system are widely reported^20,44-49^. We already mentioned the work of Cusumano and colleagues^20^, carried out in *Drosophila*, demonstrating that miR-210 overexpression in clock neurons affects large ventral lateral neurons (l-LNvs) morphology, with altered star-shaped cell bodies and aberrant PDF-positive arborisations in the optic lobes, finally resulting in visual defects. Furthermore, they found that miR-210 overexpression in all neurons or glial cells resulted in lethality^20^. Interestingly, miR-210 was identified among the miRNAs upregulated following long-term memory formation^67^ and was associated with age-related behavioural changes^68^ in the honeybee *Apis mellifera*. In other studies, miR-210 has been shown to regulate cell survival and death in neuronal cells, targeting the expression of the antiapoptotic protein Bcl-2 to induce apoptosis in neuroblastoma cells^44^ or the proapoptotic protein BNIP3 to protect against apoptosis in neural progenitor cells^45^, also inducing cell-cycle progression and terminal differentiation^46^. This apparently opposite effect, depending on cell types and conditions, should not surprise: miR-210 literature is often contradictory (*e*.*g*., on whether miR-210 action promotes^69^ or suppresses^70^ post-ischemic neurogenesis *in vivo*)^48^, and a *Janus* role of miR-210 has already been reported and characterized in cardiomyocytes subjected to hypoxia-reoxygenation stress^71^. Finally, in studies in which miR-210 expression was reduced or abolished, the decrease in miR-210 expression levels led to increased neuronal survival and improved mitochondrial function, thereby dampening proliferation in differentiating neural stem cell cultures subjected to inflammatory mediators^48^. At the same time, miR-210 loss in mouse primary hippocampal neurons cultures resulted in increased dendritic arbour density, which could be representative of altered neuronal function impacting dendritic spine and synapse formation, connectivity, and plasticity^49^. Moreover, *in vivo*, miR-210 KO mice exhibited perturbed behavioural flexibility, implying that the mechanisms governing information updating and feedback processes were affected^49^. Overall, these evidence support a role for miR-210 in neuroplasticity which might in turn represent the cause for the retinal pathological phenotype we observed and characterized in both fruit flies and mice. With this perspective in mind, we conducted an RNA-seq analysis on the brains of miR-210 KO and control flies. Earlier studies, conducted by both Weigelt^5^ and Lyu^15^, defined gene expression signatures in miR-210 KO flies using RNA-seq. However, their experiments encompassed fly heads, which included both the brain and the eyes, as well as several other tissues to a lesser extent. In contrast, our experiment was uniquely tailored to concentrate solely on the brain. We identified a substantial number of differentially expressed genes (1,053 out of 9,329 detected genes) (Table S4), with the downregulated genes predominantly enriched for the detection and transduction of light stimuli (Figure 7, Table S5). This outcome aligns with the findings of Weigelt^5^ and Lyu^15^ who studied the entire head (brain and eyes included), suggesting that the alterations observed in miR-210 KO flies extend beyond the eye and may be linked to neuronal deficiencies in signal detection and transduction. Simultaneously, we did not identify any enriched pathways related to lipid metabolism in the brains of miR-210 KO flies (Table S5). Additionally, among the upregulated genes, the few enriched pathways pertained to collagen (such as “Collagen network”, “Complex of collagen trimers”, and “Collagen type IV trimer”), thereby reinforcing the possibility of a shared upstream mechanism that underlies the retinal degeneration induced by miR-210 KO in both fruit flies and mammals. The further characterization of these models could pave the way for a complete understanding of miR-210 role in the maintenance of the proper homeostasis of the retina and of the entire visual system.

### Limitations of the study

In our research, both the miR-210 knockout fruit fly and mouse models are constitutive knockouts. To fully address the biological question concerning the role of miR-210 in the visual system, the use of tissue-specific and/or inducible miR-210 knock-out models would offer significant advantages. Concerning the fruit fly model, a potential limitation lies in the use of fly heads to examine retinal alterations; however, due to the technical challenges associated with dissecting fly retinas, employing the entire head is a common approach, as extensively documented in prior studies. As for the mouse model, our analysis was confined to the retinas of relatively young mice (aged 10-15 weeks). Investigating the retinal phenotype in older mice would be highly valuable for comprehending the long-term consequences of miR-210 depletion.

## Author contributions

Conceptualization: D.C., C.D.P.; Methodology: D.C., F.V., G.Z., P.C., M.I., F.M., C.D.P.; Formal Analysis: D.C., A.T., G.S.; Investigation: D.C., C.S.; Resources: F.V., G.Z., P.C., R.M., C.D.P.; Data Curation: D.C., G.S., C.B.; Writing – Original Draft Preparation: D.C., A.T., R.M., C.D.P.; Writing – Review & Editing Preparation: D.C., F.V., A.T., G.Z., G.S., M.I., F.M., D.T., M.M., C.B., R.M., C.D.P.; Visualization: D.C., A.T.; Supervision: M.I., F.M., D.T., M.M., C.B., R.M., C.D.P.; Project Administration: C.D.P.; Funding Acquisition: C.D.P..

## Acknowledgments

C.D.P. was supported by the following grants: PRAT 2014 - University of Padova, N. CPDA142980 and PRID 2023 - Dept. Biology, University of Padova, N. BIRD233738. F.M. was supported by the Italian Ministry of Health projects (“Ricerca Corrente”, “POS T4 CAL.HUB.RIA, cod. T4-AN-09) and by EU-NRRP M6C2 - Inv.2.1 PNRR-MAD-2022-12375790. M.I., as original generator of the miR-210 KO mouse model, was supported by the NIH/NCI RO1 grant (CA155332). We are grateful to Alessio Pollastri, Sara Boscarato, and Sofia Lombardi (University of Padova, Padova, Italy) for the technical support provided during their bachelor’s degree internships.

The authors declare no competing interests.

## Materials and Methods

### Fly strains

The following *Drosophila melanogaster* stocks were used: wDah^24^ (hereinafter referred to as wild type or WT flies), miR-210Δ^5^, GMR-GAL4 (Bloomington *Drosophila* Stock Center), UAS-miRNA-210.9^20^. Flies overexpressing miR-210 in retinal cells were obtained by crossing GMR-GAL4 and UAS-miRNA-210.9 flies. Flies were raised at 23°C under a 12/12 hours light/dark (LD) cycle and fed on a standard cornmeal-yeast agar food. The flies used for each experiment (miR-210 quantification, qRT-PCR, RNA-seq, and TEM analysis) were males of 5±1 days of age.

### Mouse models

Tissue samples from C57BL/6 miR-210 floxed mice were obtained in collaboration with Massimiliano Mazzone (Laboratory of Tumor Inflammation and Angiogenesis, Center for Cancer Biology (CCB), VIB, Leuven, Belgium)^9^ and Mircea Ivan (Department of Medicine, Indiana University, Indianapolis, Indiana, USA)^23^. Briefly, miR-210 floxed mice were crossed to Gata-1 Cre mice to constitutively delete the miR-210 locus, obtaining miR-210 KO mice. Mice were maintained under pathogen-free (SPF), temperature and humidity-controlled conditions with a 12/12 hours light/dark (LD) cycle and fed on a standard chow diet. Housing and all experimental animal procedures were approved by the Institutional Animal Care and Research Advisory Committee of the KU Leuven (Project number: 085/2020). The mice used for the behavioral analysis and the gene expression experiments (miR-210 quantification, RNA-seq, and qRT-PCR) were males of 6 and 11 weeks of age, respectively. The mice used for the confocal immunofluorescence and the transmission electron microscopy analysis were both males and females of 10-11 weeks of age. On the other hand, samples from mice overexpressing miR-210 (Doxycycline-inducible transgenic C57BL/6Ntac-*Gt(ROSA)26Sor*^*tm3720(Mir210)Tac*^ or TG-210 mice) were obtained in collaboration with the Fabio Martelli’s group (IRCCS-Policlinico San Donato, Milano, Italy)^25^. Housing and all experimental procedures complied with the Guidelines of the Italian National Institutes of Health and with the Guide for the Care and Use of Laboratory Animals (Institute of Laboratory Animal Resources, National Academy of Sciences, Bethesda, Maryland, USA) and were approved by the institutional Animal Care and Use Committee: Ministero della Salute, Direzione Generale della Sanità Animale e dei Farmaci Veterinari, authorization no. 389/2020-PR (IACUC 1038). TG-210 mice were generated by Taconic Artemis (Germany) as extensively described in Zaccagnini *et al*. (2017)^25^. Briefly, the coding region of mouse miR-210, along with 110 base pairs of its surrounding genomic sequence on either side, was introduced into the ROSA26 locus utilizing the Recombination-Mediated Cassette Exchange (RMCE) technique^26^. In order to induce miR-210 overexpression, 13 weeks old female mice were fed with food pellets containing doxycycline 2 g/kg (Mucedola srl, Milano, Italy) ad libitum for 16 days before euthanasia and samples harvesting. The eyes samples were used for miR-210 quantification and transmission electron microscopy analysis.

### Behavioral analysis of mice

For the behavioral analysis, the locomotor activity of 14 mice (7 WT and 7 miR-210 KO) were tested. Mice were monitored for 12 days under a 12/12 hours light/dark (LD) cycle and subsequently for 15 days under constant darkness (DD) to investigate daily and circadian activity. During LD cycle, lights were switched on at 7.00 a.m. (hereafter indicated as zeitgeber time and referred to as ZT0) and switched off at 7.00 p.m. (hereafter referred to as ZT12). Mice were housed in individual cages equipped with cameras and three infrared sensors, and their locomotor activity was measured as number of beam crosses and signal interruptions, collected every 1 minute. As a result, a value of zero means the absence of movement, while higher values correspond to increased locomotor activity. Given that the initial period in a new environment is typically associated with exploration rather than representative of actual locomotor activity data^27^, we focused exclusively on the data from the final 10 days under both LD and DD conditions for subsequent analysis.

### Total RNA extraction

For qRT-PCR experiments, fruit flies were frozen in liquid nitrogen and their heads were separated from the rest of the body through mechanical agitation and collected in different tubes. For different samples (muscles and fat bodies), flies were dissected in 0.1M phosphate buffered saline (PBS), then the selected tissues were transferred to ice-cold TripleXtractor reagent (GRiSP Research Solutions, Porto, Portugal). Total RNA was extracted from approximately 35 heads, 25 thoraxes (muscles), or 30 fat bodies for each sample by using the TripleXtractor reagent (GRiSP Research Solutions, Porto, Portugal) according to manufacturer’s instructions. For RNA-seq experiment, fruit fly brains were dissected in 0.1M phosphate buffered saline (PBS) and immediately transferred to ice-cold LBA-TG buffer, part of the ReliaPrep RNA Tissue Miniprep System (Promega, Madison, Wisconsin, USA), which was used to extract total RNA from approximately 30 brains for each sample according to manufacturer’s instructions. RNA concentration was measured using the NanoDrop 2000c spectrophotometer (Thermo Fisher Scientific, Waltham, Massachusetts, USA).

For both RNA-seq and qRT-PCR experiments, mice were euthanized, and the eyes enucleated and immediately put in RNAlater (Ambion, Austin, Texas, USA). For each mouse, both eyes were carefully dissected using a set of dissection tweezers and forceps (World Precision Instruments Inc., Sarasota, Florida, USA). The corneas and lenses were excised, and the retinas were isolated and pooled in 1 mL of TripleXtractor (GRiSP Research Solutions, Porto, Portugal). The entire procedure was carried out under a stereomicroscope, with the samples maintained on ice to prevent tissue degradation. Afterwards, each sample was mechanically fragmented using the IKA T10 basic ULTRA-TURRAX homogenizer (Sigma-Aldrich, St. Louis, Missouri, USA). Total RNA, including microRNAs, was isolated with the miRNeasy Mini Kit (Qiagen, Hilden, Germany) following the animal tissue protocol according to manufacturer’s instructions. RNA concentration was measured using the NanoDrop 2000c spectrophotometer (Thermo Fisher Scientific, Waltham, Massachusetts, USA) and RNA integrity was assessed by capillary electrophoresis with the RNA 6000 Nano LabChip using the Agilent Bioanalyzer 2100 (Agilent Technologies, Santa Clara, California, USA). Only samples with an RNA Integrity Number (R.I.N.) value higher than 7.0 were used for gene and miRNA expression analysis.

### Quantification of miR-210 and gene expression levels

miR-210 expression levels in both flies and mice were quantified by quantitative real-time PCR (qRT-PCR) using the miRCURY LNA miRNA PCR assay (Qiagen, Hilden, Germany). First-strand cDNA synthesis was obtained from 10 ng of total RNA by using the miRCURY LNA RT Kit (Qiagen, Hilden, Germany) following manufacturer’s instructions; 0.5 μL of UniSp6 RNA spike-in, an exogenous synthetic transcript, were added to the reaction and used as a control for monitoring the success of reverse transcription. qRT-PCR was performed in a 10 μL volume using the miRCURY LNA SYBR Green PCR Kit (Qiagen, Hilden, Germany) and the following miRCURY LNA miRNA PCR primer sets (Qiagen, Hilden, Germany): *dme-miR-210-3p* (YP02104327), *hsa-miR-210-3p* (YP00204333), *2S rRNA* (YCP2142525), *hsa-miR-16-5p* (YP00205702), and *U6 snRNA* (YP00203907). The reactions were performed in a CFX96 Touch Real-Time PCR Detection System (BioRad, Hercules, California, USA) according to the following amplification program: an initial denaturation at 95°C for 2 minutes followed by 40 cycles of denaturation at 95°C for 10 seconds and annealing/extension at 56°C for 1 minute. Each experiment included three technical replicates for each sample, and we analyzed a minimum of three independent biological replicates. The 2^-ΔΔCT^ (RQ, relative quantification) method^28^ was used to calculate miRNA relative expression levels between samples. miR-210 expression levels in flies (*dme-miR-210-3p*) and in mice (*hsa-miR-210-3p*) were normalized to *2S rRNA* and to miR-16 (*hsa-miR-16-5p*) and *U6 snRNA* respectively.

To quantify gene expression levels, we initiated the process by synthesizing first-strand cDNA from 1 μg of total RNA. We utilized the GoScript Reverse Transcriptase Kit (Promega, Madison, Wisconsin, USA) in accordance with the manufacturer’s instructions. qRT-PCR primers were designed using Primer-BLAST primer designing tool^29^ and then synthesized by Eurofins Genomics Italy srl (Milan, Italy). All used primers are listed in Table S1. The qRT-PCR reactions were conducted in a 10 μL volume using the GoTaq qPCR Master Mix chemistry (Promega, Madison, Wisconsin, USA) and a CFX384 Touch Real-Time PCR Detection System (BioRad, Hercules, California, USA). The amplification program consisted of an initial denaturation at 95°C for 2 minutes, followed by 40 cycles of denaturation at 95°C for 15 seconds and annealing/extension at 60°C for 1 minute. Each experiment included three technical replicates for each sample, and we analyzed a minimum of three independent biological replicates. The 2^-ΔΔCT^ (RQ, relative quantification) method^28^ was used to calculate relative expression levels, and *Rp49* (in flies) or *Gapdh* (in mice) were used as internal controls.

### Retinal cryosections

The mice were euthanized, the eyes were enucleated, fixed in 4% paraformaldehyde for 6 hours and washed in 0.1M phosphate buffered saline (PBS). The eyes were cryoprotected by immersion in 30% sucrose overnight, embedded in the optimum cutting temperature (OCT) compound Tissue-Tek (Qiagen, Hilden, Germany) and frozen in liquid nitrogen. Retinal cryosections of 10 μm thickness were made and collected on poly-L-lysine-coated slides through a Leica CM1850 cryostat. In order to properly compare different samples, only the retinal cryosections crossing the optic nerve were selected for the subsequent immunofluorescence staining.

### Immunofluorescence staining

5% bovine serum albumin (BSA) was used to block non-specific bindings. Cryosections were incubated overnight at 4°C with primary antibodies: polyclonal anti-GFAP (Dako, Agilent, Santa Clara, California, USA) (1:5000 in 1% BSA) and polyclonal anti-IBA-1 (Wako Pure Chemical Industries, Osaka, Japan) (1:1000 in 1% BSA). Afterwards, all cryosections were stained with secondary antibody anti-rabbit IgG conjugated to green dye Alexa Fluor 488 (Molecular Probes, Invitrogen, Carlsbad, California, USA) diluted 1:1000 in PBS, incubated at 37°C for 2 hours, and stained with bisbenzimide in order to make visible the cell nuclei.

### Confocal microscopy and images analysis

Images of immunolabeled cryosections were acquired by using a Leica TCS SP5, by setting up the same parameters for all the acquisitions. For microglia quantification, IBA-1 positive (+) cells were counted in the outer nuclear layer (ONL) through the entire section from superior to inferior. The measurement is expressed as number of IBA-1 (+) cells. To quantify GFAP levels, we focused our analysis on the central ventral retina, the region displaying structural alterations in the ONL. We employed ImageJ software to measure fluorescence intensity. Additionally, we analyzed fluorescent signals across retinal layers (ONL, OPL, INL, IPL, GCL) using ImageJ software, generating profile plots along with their corresponding grayscale intensities.

### Transmission electron microscopy (TEM)

Regarding the fruit flies, we dissected fly heads in 0.1M phosphate buffered saline (PBS) and immediately transferred them to ice-cold fixation solution containing 2.5% glutaraldehyde and fixed overnight. For the mice, the enucleated eyes were fixed in a solution composed of 2.5% glutaraldehyde and 2% paraformaldehyde in 0.1M cacodylate buffer for a duration of 4 hours. Following fixation, the retinas were carefully extracted from the eye cups and then cut into smaller pieces for further processing. Subsequently, samples were postfixed with 1% OsO4 in 0.1M sodium cacodylate buffer for 1 hour at 4°C. After three water washes, samples were dehydrated in a graded ethanol series and embedded in epoxy resin (Sigma-Aldrich, St. Louis, Missouri, USA). Ultrathin sections (60-70 nm) were obtained with a Leica Ultracut EM UC7 ultramicrotome, counterstained with uranyl acetate and lead citrate and viewed with a Tecnai G^2^ (FEI) transmission electron microscope operating at 100 kV. Images were captured with a Veleta (Olympus Soft Imaging System) digital camera.

### Triacylglycerols (TAG) quantification

Following the verification of comparable head weights between miR-210 KO and WT flies, we determined the ideal amount of undiluted tissue to be utilized for each independent sample by assessing 6 fly heads (data not shown). To induce starvation in both miR-210 KO and WT flies, we exposed them to standard conditions in tubes filled with only 1% agar in water for approximately 4 days. Triacylglycerols (TAG) quantification in the heads of both starved and non-starved miR-210 KO and WT flies was carried out using the Triglyceride Quantification Kit (Sigma-Aldrich, St. Louis, Missouri, USA), following the provided manufacturer’s instructions. Two technical replicates of each sample and at least three independent biological replicates were analyzed. At the end of the experiment, the absorbance from each well was measured at 570 nm using the Thermo Scientific Multiskan GO Microplate Spectrophotometer (Thermo Fisher Scientific, Waltham, Massachusetts, USA).

### Mouse and fly RNA-seq analyses

The RNA sequencing (RNA-seq) analysis on mouse retinas was conducted by IGA Technology Services (Udine, Italy). cDNA libraries were constructed from 100 ng of total RNA following the instructions provided by the Universal Plus Total RNA-Seq with NuQuant Kit (Tecan Genomics, Redwood City, California, USA). The workflow comprises several key steps, including the fragmentation of total RNA, cDNA synthesis using a combination of random and oligo(dT) primers, end-repair to generate blunt ends, ligation of UDI adaptors, strand selection, AnyDeplete for the removal of unwanted transcripts (such as ribosomal RNA), and PCR amplification to produce the final library. The libraries were quantified with the Qubit 2.0 Fluorometer (Invitrogen, Carlsbad, California, USA) and quality tested by Agilent 2100 Bioanalyzer High Sensitivity DNA assay. Sequencing was carried out in paired-end mode (150 bp) by using NovaSeq 6000 (Illumina, San Diego, California, USA) with a targeted sequencing depth of 80 million reads per sample. Raw data was processed with the software CASAVA v1.8.2 (Illumina, San Diego, California, USA) for both format conversion and demultiplexing. Sequence reads are available on NCBI BioProject database with the accession number PRJNA1037363.

The RNA-seq analysis conducted on fruit fly brains was performed by MicroCRIBI NGS Service (Department of Biology, University of Padova, Padova, Italy). cDNA libraries were constructed from 75 ng of total RNA by using the QuantSeq 3’ mRNA-Seq Library Prep Kit for Illumina (FWD) (Lexogen, Vienna, Austria) according to the manufacturer’s instructions. The workflow consists of first strand cDNA synthesis with oligo(dT) primers containing an Illumina-compatible sequence at the 5’ end, RNA template removal, second strand synthesis with random primers containing an Illumina-compatible linker sequence at the 5’ end, purification using magnetic beads to remove all reaction components, and PCR amplification in order to add the complete adapter sequences and to generate the final library. The libraries were quantified with the Qubit 2.0 Fluorometer (Invitrogen, Carlsbad, California, USA) and quality tested by Agilent 4150 TapeStation system (Agilent Technologies, Santa Clara, California, USA). Sequencing was carried out in single-end mode (75 bp) by using NextSeq 500 (Illumina, San Diego, California, USA) with a targeted sequencing depth of 30 million reads per sample. Base-calling was performed using RTA2 software (Illumina, San Diego, California, USA). File conversion and demultiplexing were performed using bcl2fastq software (version 2.20.0). Sequence reads are available on NCBI BioProject database with the accession number PRJNA1036442.

In both cases, raw reads were trimmed to remove adapter sequences using cutadapt (version 4.5). The abundances of all mouse and fly transcripts annotated by ENSEMBL (release 110) were estimated using the Salmon software (version 1.10.2)^30^ and then summarized at the gene level using tximport (version 1.26.0)^31^. Genes were filtered by their expression levels using the strategy described in Chen *et al*. (2016)^32^, as implemented in the edgeR package with default parameters.

In mice, a total of 13,378 genes were retained. Gene-level counts were normalized for GC-content and for unwanted variation using EDASeq (version 2.28.0) and RUVSeq (version 1.28.0; RUVg method, k=3 confounding factors)^33^. Differential expression was tested with edgeR (version 3.36.0)^34^, using a GLM model.

In fruit flies, a total of 9,329 genes were retained. Gene-level counts were normalized using the TMM method (edgeR, version 3.40.0) and differential expression was tested using a GLM model.

In both cases, genes with an adjusted p-value (FDR) < 0.05 after correction for multiple testing (Benjiamini-Hochberg method) were considered differentially expressed. Finally, in order to investigate the molecular functions of mouse and fly differentially expressed genes and the biological processes in which they were involved, a Gene Ontology (GO) functional enrichment analysis was performed using the ShinyGO tool^35^ (FDR < 0.10).

### Statistical analysis

For the behavioral analysis, we utilized the 10-day locomotor activity datasets from each mouse. Various parameters were examined using ActogramJ^36^ and subsequently compared between the two experimental groups. These parameters included the time of activity onset and offset, day-time and night-time activity, periodicity, and acrophase (which represents the time at which the peak of a rhythm occurs). Chi-square (χ^2^) periodogram analysis was used to test the presence of circadian periodicity. The daily acrophase of the locomotor activity rhythm was calculated and the average acrophase was determined by vector addition. A two-way Repeated Measures (RM) ANOVA was performed to determine significant differences.

All the results were expressed as mean ± standard error of the mean (SEM) and were obtained from at least three independent experiments. Student’s t-test, one-way ANOVA, and Bonferroni’s post-hoc test were performed to determine significant differences using the GraphPad Prism Software (version 8.0.2). p-values < 0.05 were considered statistically significant. Statistical tests and significance are described in each figure caption.

## Supplemental Information (titles and legends)

**Figure S1. Evaluation of miR-210 knock-out (KO) and overexpression (OE) in the fruit fly heads**. miR-210 expression levels in the heads of 5-day-old miR-210 KO flies (**A**) and flies overexpressing miR-210 in retinal cells (**B**) and relative controls, assessed by qRT-PCR. The results (N=3) are expressed as mean ± SEM. Student’s t-test or one-way ANOVA were performed to determine significant differences. *p-value < 0.05, **p-value < 0.01, ***p-value < 0.005.

**Figure S2. Evaluation of miR-210 expression in the heads, muscles and fat bodies of wild type fruit flies**. miR-210 expression levels in the heads, thoraxes (muscles), and fat bodies of 5-day-old wild type flies, assessed by qRT-PCR. The results (N=3) are expressed as mean ± SEM. One-way ANOVA was performed to determine significant differences. ***p-value < 0.005.

**Figure S3. Gene expression analysis in fly muscles of genes identified as differentially expressed in the heads of miR-210 KO flies**. The relative expression levels of genes implicated in the altered lipid metabolism observed in miR-210 knock-out fly heads were evaluated via qRT-PCR in the thoraxes (muscular tissue) of 5-day-old miR-210 KO flies and their respective controls. The results (N=4) are expressed as mean ± SEM. Student’s t-test was performed to determine significant differences.

**Figure S4. Triacylglycerols (TAG) quantification in the heads of starved and non-starved miR-210 KO and WT flies**. Colorimetric quantification of triacylglycerols (TAG) amount in the heads of 9-day-old starved and non-starved miR-210 KO and WT flies. The results (N=3) are expressed as mean ± SEM. One-way ANOVA was performed to determine significant differences. *p-value < 0.05, **p-value < 0.01.

**Figure S5. Assessment of miR-210 knock-out (KO) in the retinas of the mice**. miR-210 expression levels in the retinas of miR-210 KO mice and relative controls of 10-11 weeks of age, assessed by qRT-PCR. The results (N=3) are expressed as mean ± SEM. Student’s t-test was performed to determine significant differences. ***p-value < 0.005.

**Figure S6. Evaluation of miR-210 overexpression (OE) specifically in mouse retinas**. miR-210 expression levels in the retinas of mice overexpressing miR-210 and relative controls of 15 weeks of age, assessed by qRT-PCR. The results (N=3) are expressed as mean ± SEM. Student’s t-test was performed to determine significant differences. *p-value < 0.05.

**Figure S7. Transmission electron microscopy (TEM) analysis of the retina of miR-210 OE mice**. Transmission electron microscopy (TEM) images showing the photoreceptor outer segments (OS) layer in the retina of wild type (WT) and overexpressing (OE) miR-210 mice. Scale bar: 10 μm. OS = photoreceptor outer segments; IS = photoreceptor inner segments; RPE = retinal pigment epithelium. Each image is representative of at least three independent samples.

**Table S1. List of primers used for qRT-PCR**.

**Table S2. List of differentially expressed genes (DEGs) between miR-210 KO and WT mice retinas**.

**Table S3. List of biological processes (BP), cellular components (CC), and molecular functions (MF) which were significantly enriched in the comparison between miR-210 KO and WT mice retinas**.

**Table S4. List of differentially expressed genes (DEGs) between miR-210 KO and WT flies’ brains**.

**Table S5. List of biological processes (BP), cellular components (CC), and molecular functions (MF) which were significantly enriched in the comparison between miR-210 KO and WT flies’ brains**.

