## Supplementary Figures for "miR-210 is essential to retinal homeostasis in fruit flies and mice"

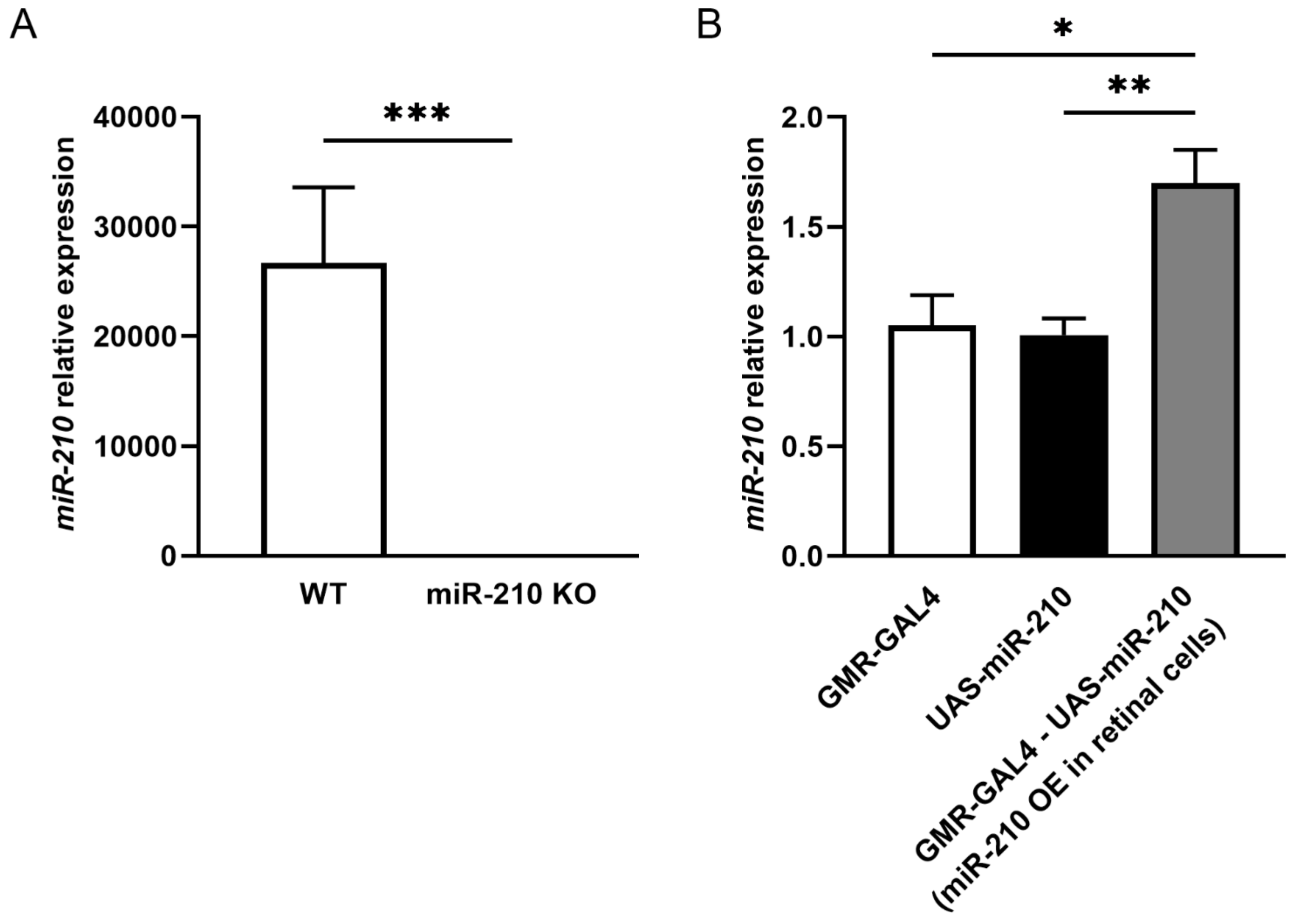


**Figure S1. Evaluation of miR-210 knock-out (KO) and overexpression (OE) in the fruit fly heads.** miR-210 expression levels in the heads of 5-day-old miR-210 KO flies (**A**) and flies overexpressing miR-210 in retinal cells (**B**) and relative controls, assessed by qRT-PCR. The results (N=3) are expressed as mean ± SEM. Student's t-test or one-way ANOVA were performed to determine significant differences. *p-value < 0.05, **p-value < 0.01, ***p-value < 0.005.

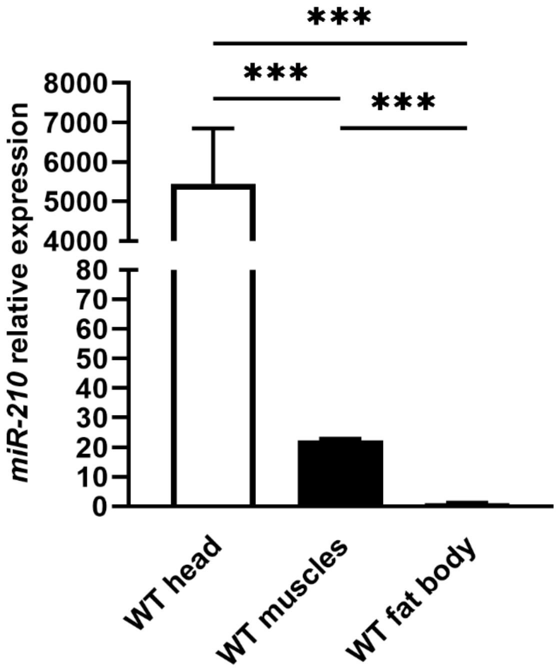


**Figure S2. Evaluation of miR-210 expression in the heads, muscles and fat bodies of wild type fruit flies.** miR-210 expression levels in the heads, thoraxes (muscles), and fat bodies of 5-day-old wild type flies, assessed by qRT-PCR. The results (N=3) are expressed as mean ± SEM. One-way ANOVA was performed to determine significant differences. ***p-value < 0.005.

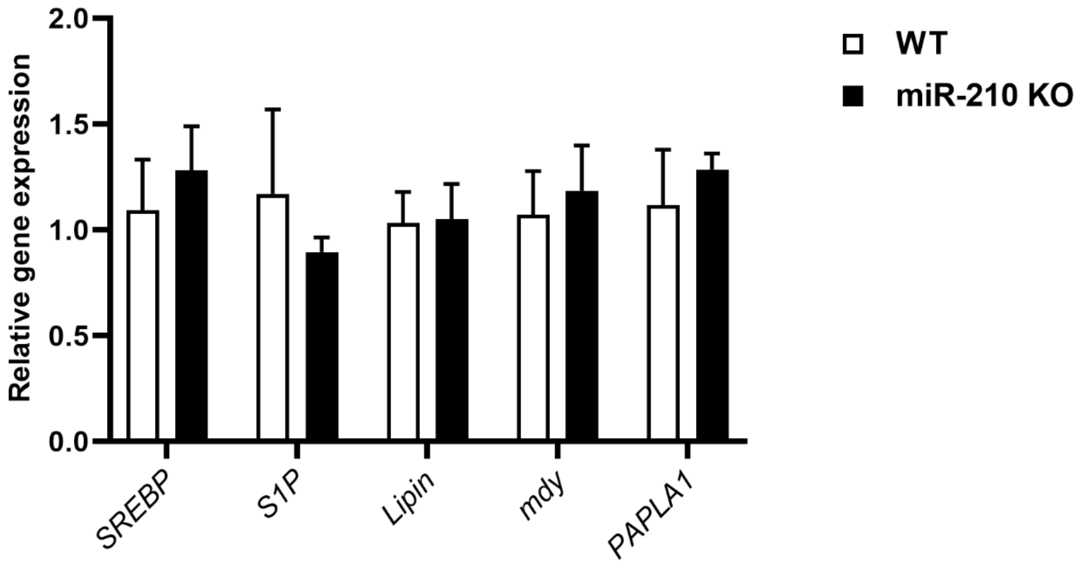


**Figure S3. Gene expression analysis in fly muscles of genes identified as differentially expressed in the heads of miR-210 KO flies.** The relative expression levels of genes implicated in the altered lipid metabolism observed in miR-210 knock-out fly heads were evaluated via qRT-PCR in the thoraxes (muscular tissue) of 5-day-old miR-210 KO flies and their respective controls. The results (N=4) are expressed as mean ± SEM. Student's t-test was performed to determine significant differences.

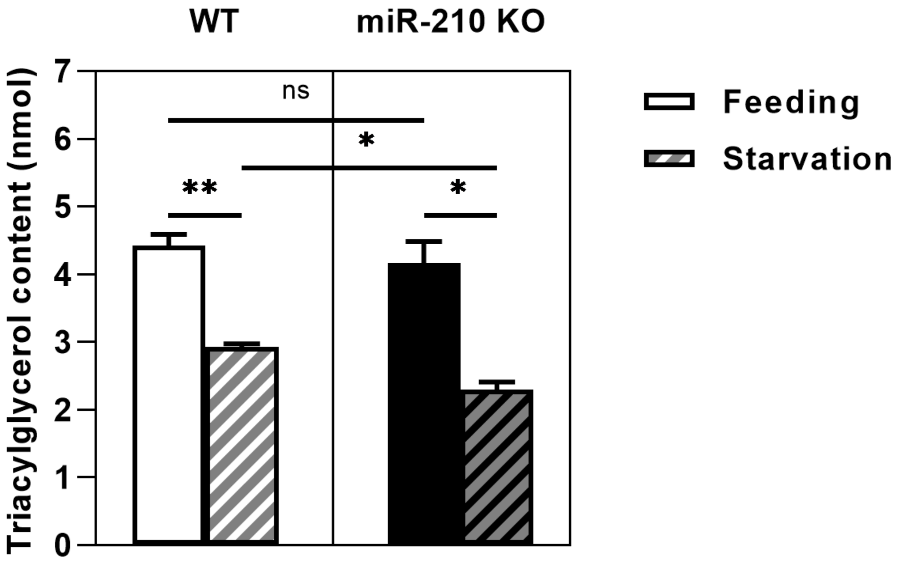


**Figure S4. Triacylglycerols (TAG) quantification in the heads of starved and non-starved miR-210 KO and WT flies.** Colorimetric quantification of triacylglycerols (TAG) amount in the heads of 9-day-old starved and non-starved miR-210 KO and WT flies. The results (N=3) are expressed as mean ± SEM. One-way ANOVA was performed to determine significant differences. *p-value < 0.05, **p-value < 0.01.

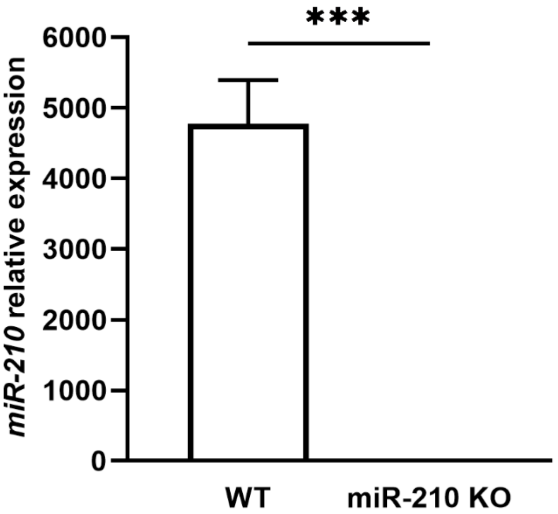


**Figure S5. Assessment of miR-210 knock-out (KO) in the retinas of the mice.** miR-210 expression levels in the retinas of miR-210 KO mice and relative controls of 10-11 weeks of age, assessed by qRT-PCR. The results (N=3) are expressed as mean ± SEM. Student's t-test was performed to determine significant differences. ***p-value < 0.005.

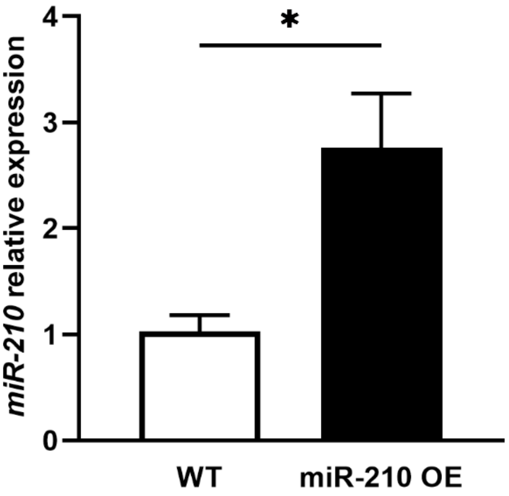


**Figure S6. Evaluation of miR-210 overexpression (OE) specifically in mouse retinas.** miR-210 expression levels in the retinas of mice overexpressing miR-210 and relative controls of 15 weeks of age, assessed by qRT-PCR. The results (N=3) are expressed as mean ± SEM. Student's t-test was performed to determine significant differences. *p-value < 0.05.

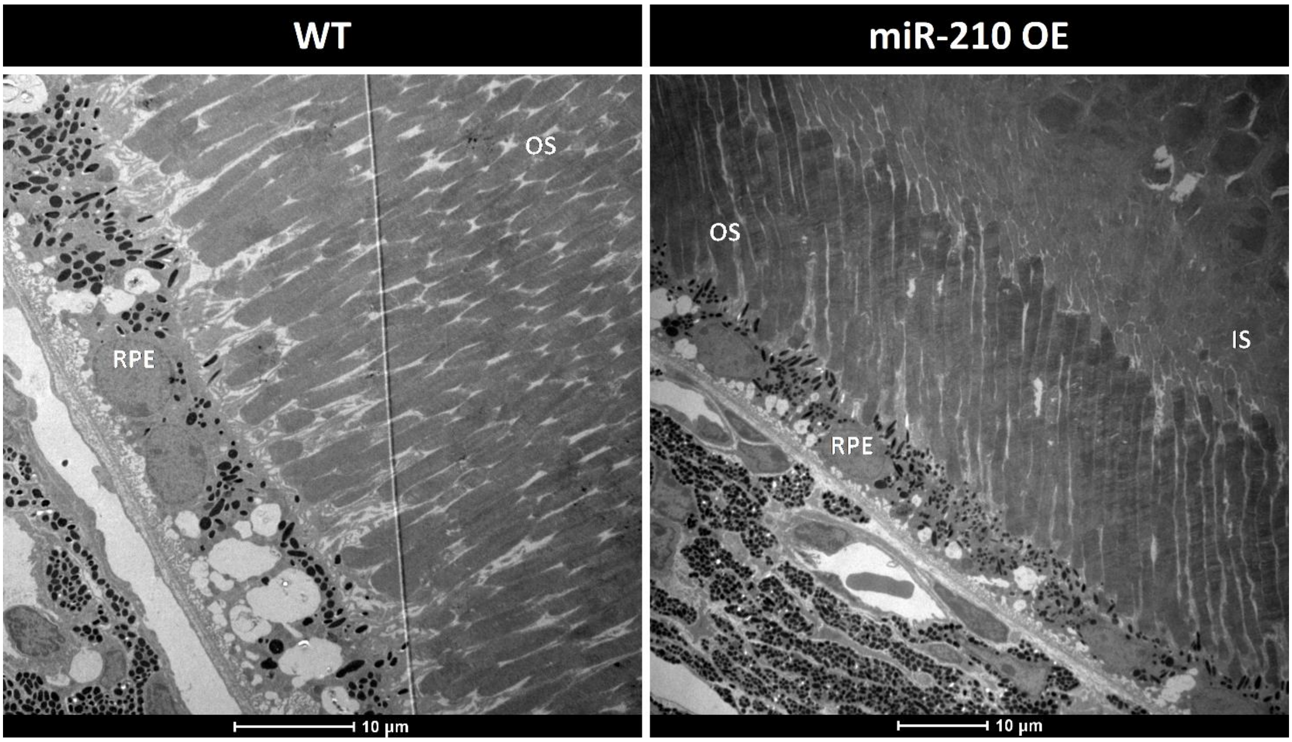


**Figure S7. Transmission electron microscopy (TEM) analysis of the retina of miR-210 OE mice.** Transmission electron microscopy (TEM) images showing the photoreceptor outer segments (OS) layer in the retina of wild type (WT) and overexpressing (OE) miR-210 mice. Scale bar: 10 µm. OS = photoreceptor outer segments; IS = photoreceptor inner segments; RPE = retinal pigment epithelium. Each image is representative of at least three independent samples.

**Table S1. List of primers used for qRT-PCR.**

**Table S2. List of differentially expressed genes (DEGs) between miR-210 KO and WT mice retinas.**

**Table S3. List of biological processes (BP), cellular components (CC), and molecular functions (MF) which were significantly enriched in the comparison between miR-210 KO and WT mice retinas.**

**Table S4. List of differentially expressed genes (DEGs) between miR-210 KO and WT flies' brains.**

**Table S5. List of biological processes (BP), cellular components (CC), and molecular functions (MF) which were significantly enriched in the comparison between miR-210 KO and WT flies' brains.**
